# The stress-inducible peroxidase *TSA*2 enables Chromosome IV duplication to be conditionally beneficial in *Saccharomyces cerevisiae*

**DOI:** 10.1101/101139

**Authors:** Robert A. Linder, John P. Greco, Fabian Seidl, Takeshi Matsui, Ian M. Ehrenreich

**Affiliations:** Molecular and Computational Biology Section, Department of Biological Sciences, University of Southern California, Los Angeles, CA 90089-2910; Department of Ecology and Evolutionary Biology, School of Biological Sciences, University of California, Irvine, Irvine, CA 92697-2525

## Abstract

Although chromosomal duplications are often deleterious, in some cases they enhance cells’ abilities to tolerate specific genetic or environmental challenges. Identifying the genes that cause particular chromosomal duplications to confer these conditionally beneficial effects can improve our understanding of the genetic and molecular mechanisms that enable certain aneuploidies to persist in cell populations and contribute to disease and evolution. Here, we perform a screen for spontaneous mutations that improve the tolerance of haploid *Saccharomyces cerevisiae* to hydrogen peroxide. Chromosome IV duplication is the most frequent mutation, as well as the only change in chromosomal copy number, seen in the screen. Using a genetic mapping strategy that involves systematically deleting segments of a duplicated chromosome, we show that the Chromosome IV duplication’s effect is largely due to the generation of a second copy of the stress-inducible cytoplasmic thioredoxin peroxidase *TSA2*. This finding is consistent with a growing literature indicating that the conditionally beneficial effects of chromosomal duplications tend to reflect the contributions of small numbers of genes that enhance tolerance to specific stresses when their copy number is increased.

**Article summary:** Changes in karyotype play an important role in evolution and health. Although these aneuploidization events are usually deleterious, in some instances they show conditionally beneficial effects by enabling cells to tolerate specific mutations or environmental stresses. The mechanisms underlying these protective effects of aneuploidization are not fully understood. To provide insights into this problem, we identify and characterize a conditionally beneficial chromosomal duplication that makes haploid yeast more tolerant to oxidative stress. We determine that the effect of the chromosomal duplication on oxidative stress tolerance is largely explained by duplication of a single stress-inducible gene.

## Introduction

Abnormalities in chromosomal copy number (or ‘aneuploidies’) often lead to cancer (Davoli *et al.* 2013; Potapova *et al.* 2013; Sheltzer 2013; Durrbaum and Storchova 2015; Laubert *et al.* 2015; Mohr *et al.* 2015; Nicholson and Cimini 2015; Pinto *et al.* 2015; Santaguida and Amon 2015; Durrbaum and Storchova 2016), developmental defects (Ottesen *et al.* 2010; Gannon *et al.* 2011; Siegel and Amon 2012; Akasaka *et al.* 2013; Bose *et al.* 2015), premature aging (Andriani *et al.* 2016; Sunshine *et al.* 2016), and other health issues in humans. In the budding yeast *Saccharomyces cerevisiae*, aneuploidies also tend to be deleterious (Torres *et al.* 2007; Yona *et al.* 2012; Potapova *et al.* 2013; Dodgson *et al.* 2016; Sunshine *et al.* 2016). However, in some cases, these aneuploidies are conditionally beneficial, as they can enable yeast to tolerate specific loss-of-function mutations or environmental stresses (Selmecki *et al.* 2009; Pavelka *et al.* 2010; Chen *et al.* 2012a; Chen *et al.* 2012b; Yona *et al.* 2012; Tan *et al.* 2013; Kaya *et al.* 2015; Liu *et al.* 2015; Meena *et al.* 2015; Selmecki *et al.* 2015; Sunshine *et al.* 2015).

An important question regarding such conditionally beneficial aneuploidies is, do their effects tend to arise due to changes in the copy numbers of one or multiple genes on the aneuploid chromosome(s)? Several studies have attempted to address this question by identifying the specific genes underlying the conditionally beneficial effects of particular chromosomal duplications (Pavelka *et al.* 2010; Chen *et al.* 2012a; Kaya *et al.* 2015; Liu *et al.* 2015). For example, Kaya *et al.* found that Chromosome XI duplication enabled S. *cerevisiae* strains lacking all eight thiol peroxidase genes to be nearly as tolerant to oxidative stress as a wild-type strain (Kaya *et al.* 2015). Experiments revealed that two genes mediated the benefit of Chromosome XI duplication: *CCP1*, a hydrogen peroxide scavenger that acts in the mitochondrial intermembrane space, and *UTH1*, a mitochondrial inner-membrane protein. In another case, Liu *et al.* showed that Chromosome VIII duplication compensates for the absence of essential nuclear pore proteins by causing overexpression of a gene that regulates cell membrane fluidity (Liu *et al.* 2015). Furthermore, Chen *et al.* demonstrated that Chr XV duplication confers resistance to the Hsp90 inhibitor radicicol by increasing dosage of *STI1*, which encodes an Hsp90 co-chaperone, and *PDR5*, which encodes a multi-drug pump (Chen *et al.* 2012a). Lastly, Pavelka *et al* showed that Chr XIII duplication confers increased resistance to the DNA-damaging agent 4-NQO by increasing dosage of *ATR1*, another multi-drug pump (Pavelka *et al.* 2010). These studies suggest that the conditional benefits of aneuploidization are typically mediated by changes in the copy numbers of a small number of genes that allow cells to cope with specific stresses.

In this paper, we explore how aneuploidies may make it possible for yeast to tolerate environmental stresses to a level beyond that achievable through genetic variation that segregates in natural populations (so-called ‘natural genetic variation’). We previously found that progeny produced by mating the lab strain BY4716 (‘BY’), the vineyard isolate RM11-1a (‘RM’), and the oak isolate YPS163 (‘YPS’) show similar maximal hydrogen peroxide tolerances despite their genetic differences (see Figure S1), suggesting the extent to which natural genetic variation can increase tolerance to this compound is limited (Linder *et al.* 2016). Here, we screen haploid segregants from these crosses for spontaneous mutations that increase hydrogen peroxide tolerance beyond the maximal levels seen for each cross in our previous study. Specifically, we take the single most tolerant F_2_ segregant that we previously identified in each of the possible pairwise crosses of the three strains and use these three segregants as the progenitors in a screen for mutations that enhance hydrogen peroxide resistance. By doing this, we obtain 37 mutants that show increased hydrogen peroxide tolerance relative to their respective progenitors.

Using whole genome sequencing, we attempt to identify the spontaneous mutations that cause increased hydrogen peroxide tolerance in the 37 mutants. Duplication of Chromosome IV (‘IV’) is the most frequent mutation, and the only aneuploidy, that we observe. Consistent with IV duplication being conditionally beneficial, we find that IV aneuploids grow worse than their progenitors in the absence of hydrogen peroxide and that the benefit of IV disomy occurs on agar plates but not in liquid media. Following these discoveries, we attempt to determine the genetic basis of this chromosomal duplication’s effect using chromosome- and gene-scale genetic engineering. Employing these techniques, we identify a single gene, the stress-inducible cytoplasmic thioredoxin peroxidase *TSA2*, which accounts for the effect of IV duplication on hydrogen peroxide tolerance. Our findings illustrate how aneuploidies may enable cells to tolerate stress at a level beyond that achievable through natural genetic variation and provide further support that the conditionally beneficial effects of aneuploidies tend to have a simple genetic basis.

## Materials and Methods

### Screen for increased hydrogen peroxide tolerance

Progenitor strains were produced during our past work on the BYxRM, BYxYPS, and RMxYPS crosses (Ehrenreich *et al.* 2012; Linder *et al.* 2016) and are described in more detail in (Linder *et al.* 2016). Each progenitor strain was streaked onto yeast extract-peptone-dextrose (YPD) plates and incubated for two days at 30°C. To maximize biological independence among different mutations obtained from the screen, 24 different colonies per progenitor were each inoculated into 800 μl of YPD broth. These cultures were outgrown for two days at 30°C with shaking at 200 RPM. 20 μl from each culture were then diluted using 80 μl of sterile water and spread onto YPD plates containing a range of hydrogen peroxide doses. These plates were incubated at 30°C for four to six days, so that slow growing mutants would have enough time to form visible colonies. Glycerol freezer stocks were then made for mutants that formed visible colonies at doses at least 1 mM higher than the minimum inhibitory concentration (MIC) of their progenitor and stored at −80°C.

All mutants were phenotyped side-by-side with their respective progenitors across a broad range of hydrogen peroxide doses to confirm their increased tolerance. Mutants that grew at doses at least 0.5 mM higher than their progenitor were saved for downstream analysis. These included 14 BYxRM-, 9 RMxYPS-, and 14 BYxYPS-derived mutants.

### Genome sequencing of mutants

Archived mutants were inoculated into YPD liquid and outgrown for two days at 30°C. For each mutant, DNA was extracted using the Qiagen DNeasy kit and a whole genome library was prepared using the Illumina Nextera kit. Each library was tagged with a unique DNA barcode to enable multiplexing. Sequencing was done on an Illumina NextSeq 500 instrument at the USC Epigenome Center. Reads were then demultiplexed using custom Python scripts.

Progenitor strains were sequenced to an average of 116X coverage to generate strain-specific reference genomes, which short read data from the mutants could then be mapped against and used to identify *de novo* mutations. The Burrows-Wheeler Aligner (BWA-MEM) (Li and Durbin 2009) was used to map reads from the progenitors to the BY genome. The parameters used for alignment were: ‘bwa –mem –t 6 ref.fsa read1 .f1 read2.fq > output.sam’. To remove duplicate reads, the rmdup command was used in SAMtools (Li *et al.* 2009). In order to generate Mpileup files, SAMtools was then implemented with the command ‘samtools mpileup –f ref.fsa read.rmdp.srt.bam > output.mp’. To generate a reference genome for a given progenitor, we identified genetic differences between the progenitor and BY, integrated these differences into the BY genome, and then re-mapped reads. This process was repeated for up to 10 cycles and the output genome was then used as the reference for mapping reads from mutants derived from that progenitor.

Mutant strains were sequenced to an average of 130X coverage (see Note S1). Reads from mutant strains were aligned to the progenitor reference genomes using BWA-MEM and the same parameters described above, followed by the generation of mpileup files with SAMtools. Mutations were identified as differences from the corresponding progenitor that were seen in at least 90% of the reads at a particular genomic site. Custom Python scripts were used to identify these mutations, as well as to calculate per site or per genomic window sequencing coverage.

### Gene Ontology enrichment analysis

GO analysis was carried out on the *Saccharomyces* Genome Database website (http://www.yeastgenome.org/cgi-bin/GO/goTermFinder.pl) using GO Term Finder version 0.83 with the molecular function category selected. All genes listed in Tables S1 and S2 were included in the analysis.

### PCR-mediated chromosomal deletion

Similar to (Sugiyama *et al.* 2008), PCR-mediated chromosomal deletion (PCD) was implemented using constructs with three segments (in this order): 300 to 600 bp of sequence homologous to the desired integration site on IV, a *kanMX* cassette, and a synthetic telomere seed sequence consisting of six repeats of the motif 5’-CCCCAA-3’ (Figure 1). To generate this construct, the region containing the integration site was PCR amplified from genomic DNA using a reverse primer that was tailed with 30 bases of sequence identity to the *kanMX* construct. At the same time, *kanMX* was PCR amplified from a plasmid using a forward primer with 30 bases of sequence identity to the integration site and a reverse primer containing the synthetic telomere seed sequence.

**Figure 1.**
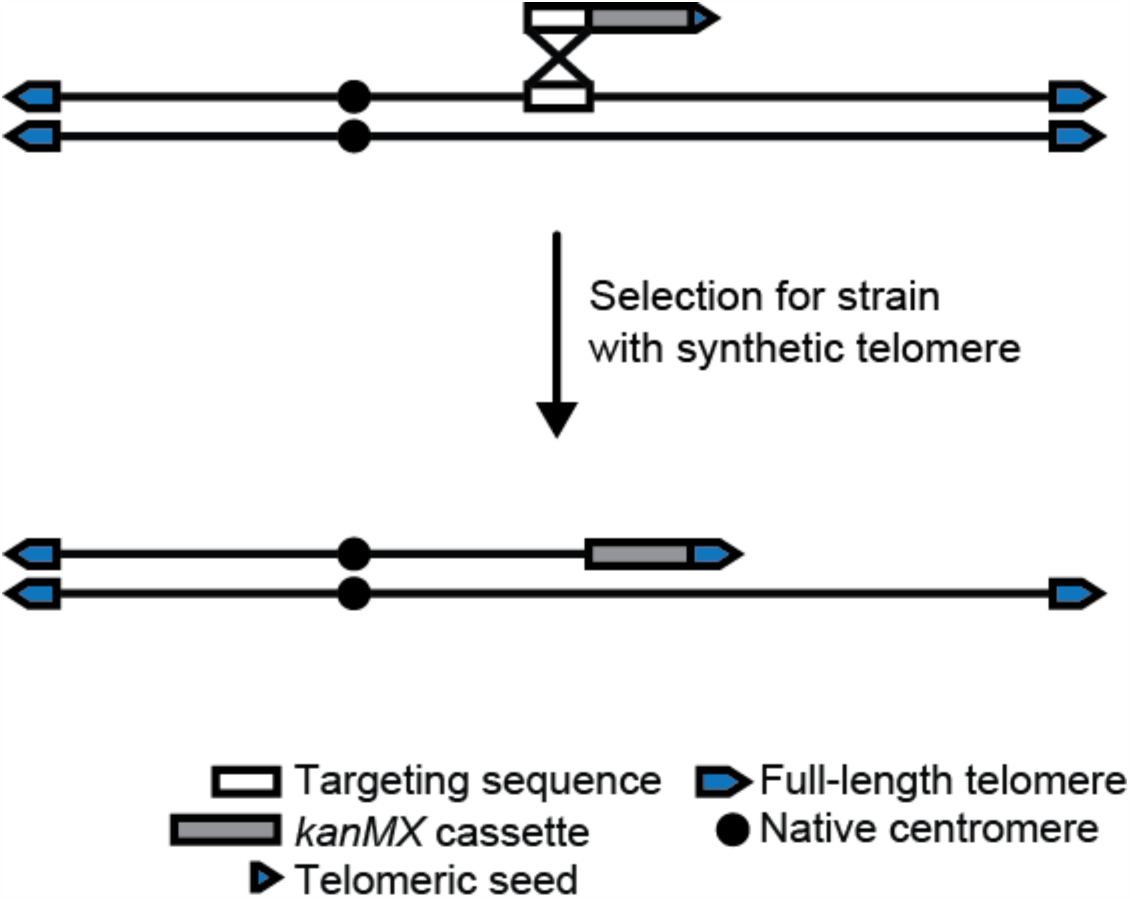
PCR-mediated chromosome deletion (PCD) was used to eliminate duplicated chromosomal segments from a BYxRM-derived aneuploid that possessed two complete copies of Chromosome IV. As described in the Methods, each PCD construct consisted of a sequence identical to a particular site on the chromosome, a *kanMX* marker providing resistance to G418, and a synthetic telomere seed sequence.

Integration site and *kanMX*/synthetic telomere seed sequences were joined using overlap fusion PCR (Sugiyama *et al.* 2005). This was done by mixing the two products in equimolar fractions in combination with the forward primer for the integration site and the reverse primer for the *kanMX*/synthetic telomere seed sequence, and conducting PCR. The cycling parameters for overlap fusion PCR were as follows: initial denaturation at 98°C for 3 minutes, followed by 30 cycles of 98°C for 30 seconds, 63°C for 30 seconds, and 72°C for 1.5 minutes. The final step was a 5 minute extension at 72°C. Throughout this process, all PCR steps were implemented using NEB High-Fidelity Phusion polymerase and all PCR products were purified using either Qiagen QIAquick Gel Extraction or QIAquick PCR Purification kits.

A standard lithium acetate-based technique was used to transform PCD constructs into cells, with about 5 μg of construct employed (Gietz and Schiestl 2007). Transformants were recovered using selection for G418 resistance on YPD plates, verified by colony PCR or genome sequencing, and archived at −80°C in glycerol solution. Primers used to construct PCD products are listed in Table S3.

### Deletion of individual genes

Targeted gene deletions were performed using the *kanMX* cassette. Constructs for deleting *ECM11, ADA2, UTP6, NHX1*, and *GUK1* were generated by amplifying the *kanMX* cassette from a plasmid using tailed primers. Each tail contained 30 to 60 bases of sequence homology to the target gene's flanking regions. Specifically, the forward and reverse primers were designed to have tails identical to the region immediately upstream of the translation start site and downstream of the stop codon, respectively.

Regarding deletions of *YDR455C, PPN1, TOM1, TSA2, YHP1, RPS18A, and RPS17B*, transformations using knockout cassettes tailed with only 60 to 120 bp of total homology to the targeted region were relatively inefficient. To increase the efficiency of these knockouts, gene deletion products were generated in a manner analogous to making the PCD constructs described above. 300 to 600 bp of sequence targeting a region just upstream of the translation start site was fused to the *kanMX* cassette, with the caveat that the reverse primer for *kanMX* amplification was tailed with 30 to 60 bases of homology to a region just downstream of the gene being deleted. Additionally, several hundred bp of sequence up- and downstream of *TSA2* were deleted by the same method, using a modified targeting sequence and downstream homology tail.

The same lithium acetate-based transformation methods described for PCD were used for all individual gene knockouts. Transformants were verified by colony PCR. Primers used for individual gene knockouts are listed in Table S4.

### Phenotyping of transformants

Transformants were outgrown for two days in 800 μL of YPD broth incubated at 30°C with shaking. As controls for batch effects, each time one or more transformants were phenotyped, their euploid and aneuploid progenitors were also examined. After the liquid outgrowth step, strains were pinned onto YPD plates supplemented with a range of hydrogen peroxide doses. Alternatively, in some experiments, strains were then transferred to liquid media supplemented with a range of doses of hydrogen peroxide, after which they were pinned onto YPD plates. Each experiment was done in biological triplicate. Plates were incubated for three days at 30°C and then imaged on a GelDoc imaging device using a 0.5 second exposure time. MIC was determined as the lowest hydrogen peroxide dose at which a given strain could not grow.

## Results and Discussion

### Screen for spontaneous mutations that increase hydrogen peroxide tolerance

24 independent cultures of each of the three haploid progenitor strains were examined after two days (20 generations) of outgrowth using selection on agar plates supplemented with hydrogen peroxide (Methods). All mutants (37 total) that exhibited an increase in minimum inhibitory concentration (MIC) at least 0.5 mM higher than their corresponding progenitor were analysed further (see Figure S2A-C; Methods). BYxRM-, RMxYPS-, and BYxYPS-derived mutants were, on average, 2.3 mM (32%), 2.1 mM (25%), and 0.6 mM (7%) more tolerant than their progenitor, respectively (see Figure S2A-C).

### The most frequently identified mutation is a chromosomal duplication

Analysis of genome-wide sequencing coverage indicated that 43% of the mutants carried two complete copies of IV (Figure 2). No other aneuploidies were detected. The disomy was common among the BYxRM-(79%) and RMxYPS-derived (45%) mutants, but was absent from the BYxYPS-derived mutants (Figure 2B). Given that the BYxRM- and RMxYPS-derived mutants also showed higher average gains in tolerance, this finding is consistent with duplication of IV conferring a sizable increase in tolerance (see Figure S2A-C).

**Figure 2.**
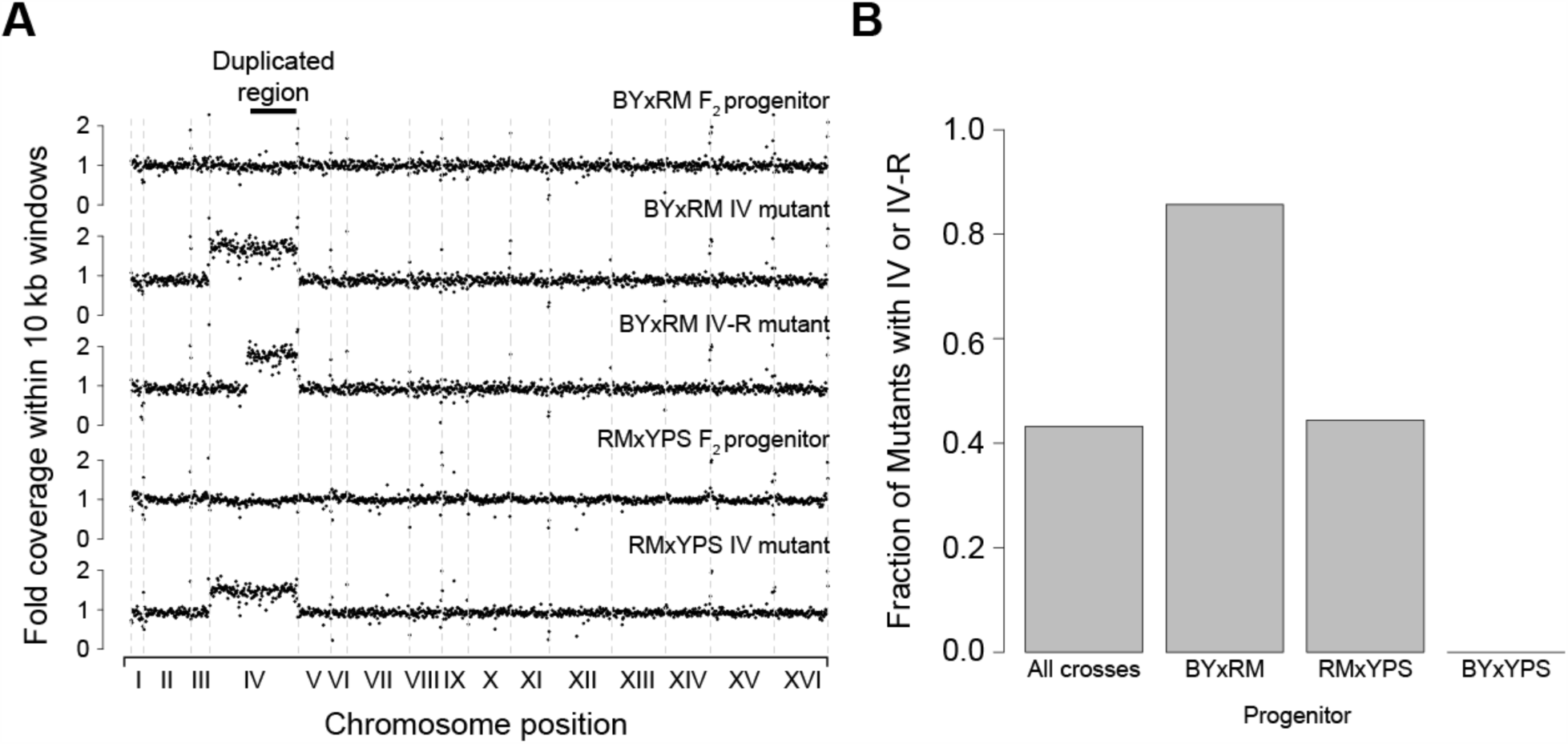
Chromosome IV duplication is the most frequent mutational event in a screen for spontaneous hydrogen peroxide resistance mutations. (**A**) Genome-wide coverage plots are provided for the BYxRM and RMxYPS F_2_ progenitors, as well as representative aneuploid or segmental duplication mutants derived from them. (**B**) Bar plots show the fraction of sequenced mutants that were disomic for the right arm of Chromosome IV both across the entire screen and by individual progenitor.

A single segmental duplication was also detected; this was found in a BYxRM-derived mutant that possessed two copies of only the right arm of IV (‘IV-R’; Figure 2A). The segmental duplication spanned approximately 890 kb (58% of IV). Including this mutant, 86% of the BYxRM-derived mutants were disomic for IV-R.

Additionally, 39 unique point mutations were identified among the mutants (see Table S1; Table S2). Most of these point mutations were non-synonymous (22) or noncoding (5) changes, some of which occurred in genes that are known to affect hydrogen peroxide tolerance, including a SUMO E3 ligase involved in DNA repair (*MMS21*), an activator of cytochrome oxidase 1 translation (*PET309*), a subunit of cytochrome c oxidase (*COX1*), and a negative regulator of Ras-cAMP-PKA signalling (*GPB2*; see Table S1; Table S2). At False Discovery Rates of 0.17 or lower, no specific Gene Ontology enrichments were seen among the genes harboring point mutations (Methods). These results, although consistent with our past finding that genetic perturbation of many different cellular processes can influence hydrogen peroxide tolerance (Linder *et al.* 2016), could reflect the inclusion of passenger mutations that do not play causal roles in hydrogen peroxide tolerance in our analysis.

Duplication of IV-R was the most frequently detected mutation in our screen. However, strains carrying point mutations were found among the BYxRM- and RMxYPS-derived mutants that were as hydrogen peroxide tolerant as strains containing the IV-R duplication (see Figure S2A and B). This suggests that duplication of IV-R may have occurred more often in our screen than point mutations because spontaneous aneuploidies occur more frequently than spontaneous point mutations in certain genetic backgrounds (Kaya *et al.* 2015; Liu *et al.* 2015). Furthermore, similar to past reports that chromosomal duplications are usually deleterious (Pavelka *et al.* 2010; Sunshine *et al.* 2016), IV-R duplication both reduced overall growth on rich media as compared to the euploid progenitor (Figure 3A; Figure S3) and only conferred a benefit when hydrogen peroxide exposure occurred on agar plates (Figure 3B and C). This implies that duplication of IV-R is conditionally beneficial.

**Figure 3.**
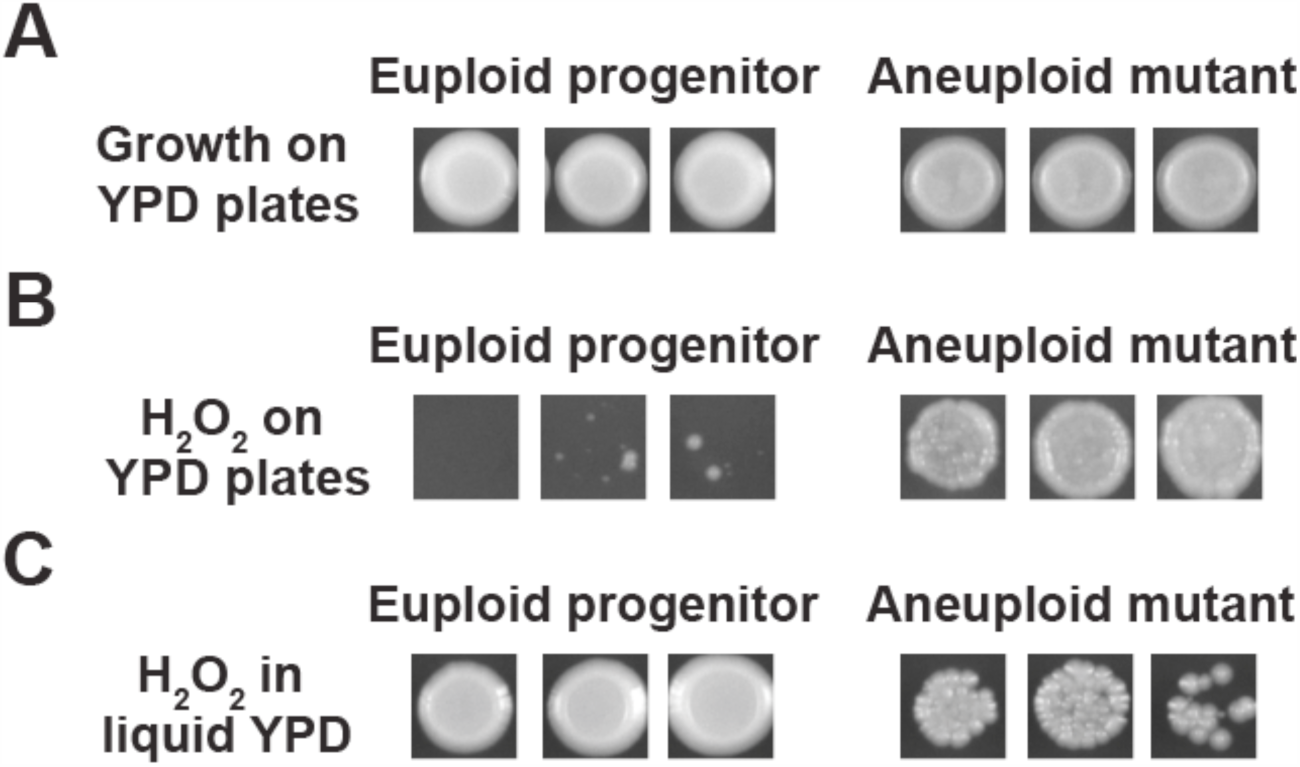
Duplication of IV is conditionally beneficial. In **(A)**, euploid replicates show superior overall growth on agar plates with rich media as compared to replicates of a strain fully disomic for IV. However, when replicates of these strains are grown on agar plates supplemented with 7.5mM hydrogen peroxide **(B)**, individuals disomic for IV show significantly improved growth as compared to euploid individuals. Additionally, when replicates of both strains are exposed to 40mM of hydrogen peroxide in liquid culture and subsequently pinned onto agar plates with rich media **(C)**, euploid individuals show significantly increased growth as compared to individuals disomic for IV. In **(A)**, representative individuals are depicted. For **(A)** and **(B)**, individuals were pinned onto agar plates with or without hydrogen peroxide supplementation and imaged after three days (Methods). In **(C)**, individuals were exposed to hydrogen peroxide in liquid media for three days, after which they were pinned onto agar plates with rich media. Images were taken after three days of growth (Methods).

### Identification of a single region responsible for most of the effect of IV-R duplication

To map the conditional growth benefit of IV-R duplication to specific genes, we adapted a technique known as PCR-mediated chromosomal deletion (‘PCD’), which involves eliminating segments of a chromosome that are distal to a centromere by inserting a drug resistance cassette linked to a synthetic telomere seed sequence (Figure 1; Methods) (Sugiyama *et al.* 2005; Sugiyama *et al.* 2008; Kaboli *et al.* 2016). Colony PCR was used both to confirm correct placement of the deletion cassette as well as to verify that a single copy of the deleted region remained (Methods).

We first used PCD to delete the right half of IV-R from a BYxRM-derived aneuploid (Figure 4A; Methods). This genetic change, which was confirmed by whole genome sequencing (see Figure S4; Methods), caused a reversion to the hydrogen peroxide tolerance exhibited by the euploid BYxRM progenitor prior to the screen (see Figure S5A). This result indicates that the deleted chromosomal segment is required for the aneuploidy's effect.

**Figure 4.**
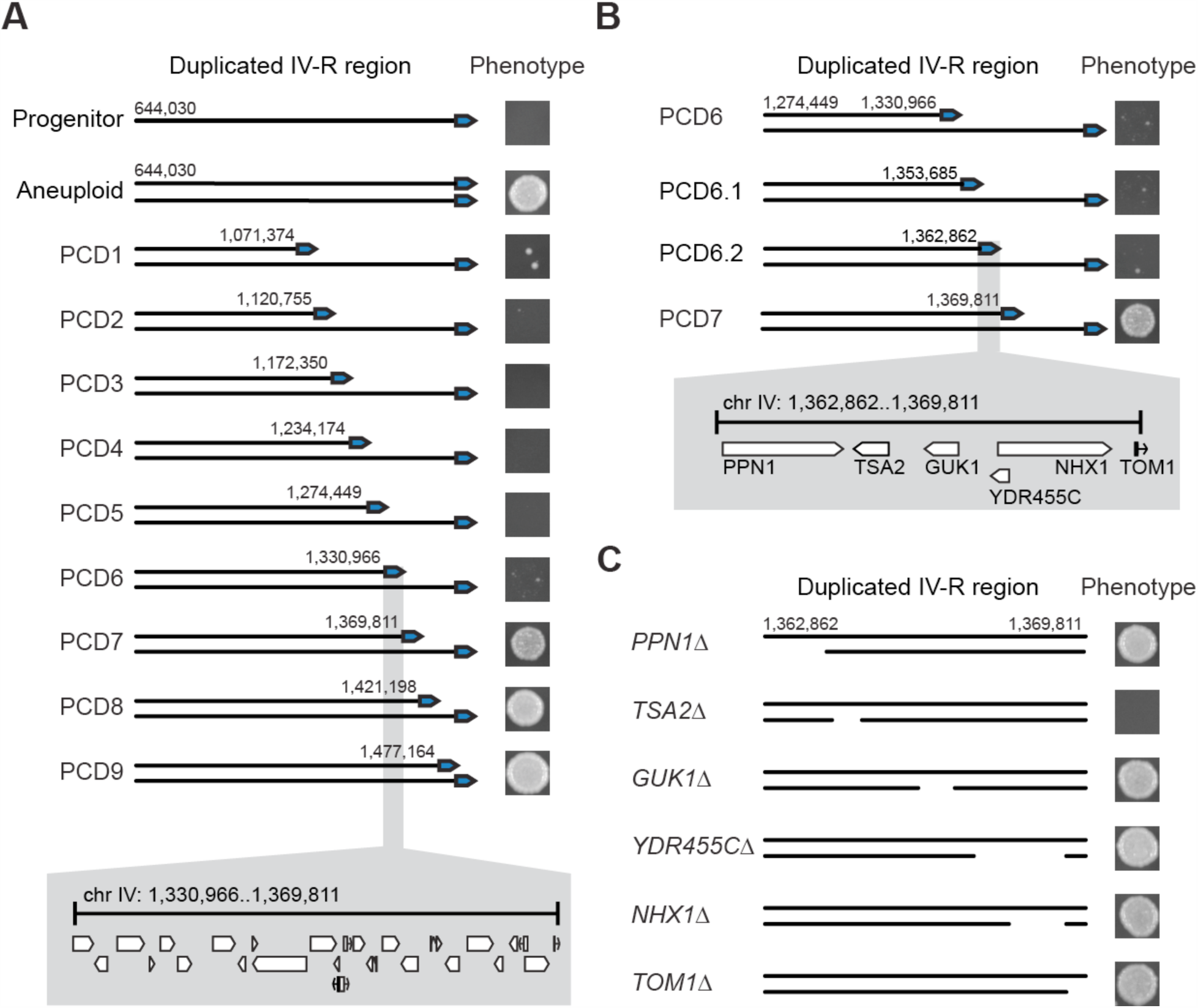
Genetic dissection of the Chromosome IV duplication's effect on hydrogen peroxide tolerance. (**A**) PCDs were staggered nearly every 50 kb along IV-R in a BYxRM-derived aneuploid. Phenotyping of partially aneuploid strains generated by PCD identified a single genomic interval with a large effect on hydrogen peroxide tolerance when strains were examined at a dose of 7.5 mM. This approximately 40 kb region contains 23 genes and 3 dubious ORFs. (**B**) Two additional PCD strains were generated within the previously mentioned window and examined at 7.5 mM. This narrowed the interval to 7 kb that contained five genes and a dubious ORF. **(C)** Individual gene deletions revealed that *TSA2* is largely responsible for the increase in tolerance conferred by duplication of Chromosome IV-R. In **A-B**, representative images of colonies grown at 7.5mM of hydrogen peroxide are shown next to their associated PCD strains; numbers adjacent to the telomere indicate the starting position of each PCD on Chromosome IV. In **C**, representative images of colonies grown at 7.5mM of hydrogen peroxide are again shown next to their associated individual gene deletion strains, with deleted regions indicated by gaps on the duplicated chromosome.

We next generated a panel of PCD strains, with large-scale deletions staggered, on average, every 50.7 kb along the latter half of IV-R. This led to the identification of a single 40 kb region that recapitulates most of the effect of the IV-R disomy (Figure 4A; Figure S5A). It is important to note that, although chromosome-scale deletions of this 40 kb region appeared to phenocopy the progenitor at certain hydrogen peroxide doses (Figure 4A and B), the average MICs of these strains were higher than that of their progenitor (Figure S5A and B). Possible explanations for this result include the presence of one or more additional point mutations that contribute to hydrogen peroxide tolerance in the mutant, an additional gene or regulatory element on IV-R whose dosage contributes to tolerance, and non-linear relationships between DNA content and hydrogen peroxide tolerance, as the chromosome-scale deletion strains remain aneuploid at a large portion of IV (see Note S2).

### Duplication of TSA2 mediates the conditionally beneficial effect of the aneuploidy

To further resolve this window, we performed two additional PCD transformations, which fine-mapped the causal interval to roughly 7 kb, spanning positions 1,362,862 bp to 1,369,812 bp (Figure 4B; Figure S5B). This interval contains five genes—the polyphosphatase *PPN1*, the cytoplasmic thioredoxin peroxidase *TSA2*, the guanylate kinase *GUK1*, the ion exchanger *NHX1*, and the E3 ubiquitin ligase *TOM1*—as well as a dubious ORF *(YDR455C).* We used standard techniques to individually delete each of these genes from a BYxRM-derived aneuploid, again using colony PCR to verify both correct placement of the deletion cassette and that a single copy of the gene remained (Methods). Also, because telomeres can influence the transcription of genes more than 20 kb away (Gottschling *et al.* 1990; Aparicio and Gottschling 1994), we deleted the six genes upstream of *PPN1* (Figure 4B; Figure S6).

The only gene deletion that showed a phenotypic effect was *TSA2*, which encodes a cytoplasmic thioredoxin peroxidase (Figure 4B and C; Figure S6) (Gasch *et al.* 2000; Park *et al.* 2000; Wong *et al.* 2002; Munhoz and Netto 2004; Ogusucu *et al.* 2007; Nielsen *et al.* 2016). Previous work (Wong *et al.* 2002) has shown that deleting *TSA2* leads to a decrease in hydrogen peroxide tolerance. Loss of the *TSA2* coding region eliminated the majority of the aneuploidy’s effect (Figure 4C; Figure S6), proving a causal role for *TSA2* in the conditional benefit conferred by IV-R duplication. Knockout of *TSA2* in other mutants, including a fully disomic BYxRM mutant, a partially disomic BYxRM mutant, and a fully disomic RMxYPS mutant, confirmed that the effect of *TSA2* was reproducible across different aneuploid individuals recovered from our screen (Figure S7; see Note S3).

## Conclusion

Previously, we found that the maximal hydrogen peroxide tolerances of segregants from the BYxRM, BYxYPS, and RMxYPS crosses are comparable (see Figure S1) (Linder *et al.* 2016), suggesting that the level of tolerance achievable through natural genetic variation may be constrained to some degree. To surpass the levels of tolerance seen in our prior study, we conducted a screen for spontaneous mutations that confer higher hydrogen peroxide tolerance than we previously observed. We employed segregants that had maximal tolerances as the progenitors in our mutagenesis screen because these individuals carry combinations of naturally occurring alleles that lead to resistance and we wanted to find mutations that provide even greater tolerance than these allele combinations.

Duplication of IV-R was the most common mutation in our screen. As with other chromosomal duplications (Pavelka *et al.* 2010; Sunshine *et al.* 2015), the beneficial effect of the IV-R disomy is conditional: it depends on both the presence of hydrogen peroxide and exposure to hydrogen peroxide on agar plates, and may be genetic background-dependent. Regarding this latter point, the present data cannot differentiate whether the progenitors in our screen varied in their genome stabilities or in their abilities to beneficially utilize the IV-R disomy. Assessment of genotype at *TSA2* indicates that pre-existing variation at this locus probably does not explain why we observed IV aneuploids in the BYxRM and RMxYPS crosses, but not the BYxYPS cross (see Note S4). Regardless, our work clearly supports the idea that myriad contextual factors influence the potential for IV-R duplication to confer beneficial effects.

Using chromosome- and gene-scale deletions, we determined that increased copy number of a single detoxifying gene, *TSA2*, explains the majority of the benefit conferred by duplication of IV-R. *TSA2* is unique among budding yeast’s cytoplasmic thioredoxin peroxidases, as it is the only one that shows markedly increased activation in response to hydrogen peroxide (Gasch *et al.* 2000; Park *et al.* 2000; Wong *et al.* 2002; Munhoz and Netto 2004; Ogusucu *et al.* 2007; Nielsen *et al.* 2016). *TSA2’s* distinctive environmental responsiveness may help to explain not only why duplication of IV-R is conditionally beneficial, but also why this was the only aneuploidy recovered in our screen. Our results are consistent with recent studies from other groups showing that typically a small number of genes, usually one or two, mediate the conditionally beneficial effects of aneuploidies (Pavelka *et al.* 2010; Chen *et al.* 2012a; Kaya *et al.* 2015; Liu *et al.* 2015).

In summary, our work speaks to challenges in enhancing particular traits using natural genetic variation, spontaneous mutations, or a combination of the two. Indeed, the maximal trait values achievable through natural genetic variation may be limited, both due to the specific alleles present in a population and due to features of a system that prevent extreme phenotypic levels from occurring (Kryazhimskiy *et al.* 2015). Trying to overcome these limits may be achievable using spontaneous mutations, but our work suggests the most likely mutational event to underlie such phenotypic increases is chromosomal duplication. Such duplications have diminished utility because they are easily lost, meaning that phenotypic gains can quickly revert (Berman 2016). This is why some have argued, within the context of evolution, that aneuploidies may be a temporary state that facilitates the acquisition of other mutations that provide a more permanent solution to stressful conditions (Yona *et al.* 2012). It may therefore be necessary to further evolve strains with a conditionally beneficial duplication(s) to enable the acquisition of stable genetic changes that can more permanently enhance a particular trait.

## Acknowledgments

We thank Jonathan Lee, Takeshi Matsui, Martin Mullis, and Rachel Schell for reviewing a draft of this manuscript. This work was supported by grants from the National Institutes of Health (R01GM110255), National Science Foundation (MCB1330874), and Alfred P. Sloan Foundation to I.M.E.

